# Some lessons and perspectives for applications of stochastic models in biological and cancer research

**DOI:** 10.1101/270215

**Authors:** Alan U. Sabino, Miguel F. S. Vasconcelos, Misaki Y. Sittoni, Willian W. Lautenschläger, Alexandre S. Queiroga, Mauro C. C. de Morais, Alexandre F. Ramos

## Abstract

Randomness is an unavoidable feature of inner cellular environment and its effects propagate to higher levels of living matter organization such as cells, tissues, and organisms. Approaching those systems experimentally to understand their dynamics is a complex task because of the plethora of compounds interacting in a web that combines intra and inter level elements such that a coordinate behavior come up. Such a characteristic points to the necessity of establishing principles that help on the description, categorization, classification, and the prediction of the behavior of biological systems. The theoretical machinery already available, or the ones to be discovered motivated by biological problems, can play an important role on that quest. Here we exemplify the applicability of theoretical tools by discussing some biological problems that we have approached mathematically: fluctuations in gene expression and cell proliferation in the context of loss of contact inhibition. We discuss the methods that we have employed aiming to provide the reader with a phenomenological, biologically motivated, perspective of the use of theoretical methods. Furthermore, we discuss some of our conclusions after employing our approach and some research perspectives that they motivate.

## I. INTRODUCTION

Description of Physical phenomena by means of theoretical methods has motivated building a rich machinery spreading from general relativity (description of matter behavior at macroscopic scale) to quantum mechanics (description of matter behavior at microscopic scale), from electromagnetism (description of electric charges, magnetic dipoles and light related phenomena) to condensed matter theory (microscopic description of solid state systems). Those tools have enabled the capability of controlling and designing specific experiments which outcomes are, in general, predicted within specific error ranges or to develop new technologies derived from that knowledge. Fortunately, theories have a scope of applicability which means they do not explain all observed data related to a given phenomenon. In general, that motivates elaborating new theories that may raise additional predictions, *i.e.*, additional verifiable hypothesis. For example, differently of Newtoniang gravity, general relativity succesfully predicts precession of the perihelion of Mercury or light bending by Sun. Furthermore, experiments aiming to investigate different manifestations of a phenomenon would require the development of specific theoretical or technological tools. For instance, one may consider the use of tensor calculus in general relativity instead of vector calculus of Newtonian gravitation, or the high precision instruments required for detection of gravitational waves. Biology, on the other hand, has followed a different historical trajectory with a predominant use of experimental methods. Biologists also rely on qualitative models to aid on the construction of a static picture of biological phenomena. Such an approach has relevant scientific and technologic implications. For example, one may consider the establishment of evolution theory - a key paradigm of modern science - or the capability of controlling biological phenomena at molecular level - as it is the case of human insulin production. However that strategy has a clear limit if one is interested on the dynamics resulting of the interaction of a plethora of compounds at different levels of living matter organization, such as organisms, tissues, cells and molecules. Additionally, interactions among elements of different levels give birth to a highly complex picture which description will demand the use of all machinery available in the scientific toolbox. That includes the use of mathematical methods not only as a crunching number technique but also as a strategy for formulating new principles that might be useful on description of biological phenomena, on testing hypothesis not accessible experimentally, and, for the case of successful theories, predicting the outcomes of different experimental designs, or guide the development of new technologies.

In this mini review we approach the biological usefulness of quantitative techniques in investigation of biological phenomena. We consider the application of stochastic methods to describe phenomena happening at molecular and cellular levels. The first topic will be reviewed within the scope of a stochastic model for binary regulation of expression of a gene that is either self-repressed or externally regulated. The second topic is approached in terms of a stochastic model aiming to quantify the role of contact inhibition in a co-culture *in vitro* experiment combining keratinocytes and melanoma cells. Our molecular level investigations have enabled us to understand the characteristics of chemical kinetics of ON and OFF switch of a gene that leads self-repression to be the mechanisms of noise reduction on amounts of expressed proteins of a given gene [1]. Furthermore, we used this model to approach noise in development of *D. melanogaster* embryos. That helped us to understand how the externally regulating gene enables mRNA production and spatio-temporal patterning with proper precision during development [2]. At cellular level, it was shown that cell proliferation mechanism denoted as contact inhibition can be quantified as an exclusion diameter between cells. That mechanism enables the formation of clusters of melanoma in co-culture with keratinocytes [3]. Furhtermore, that model predicts that melanoma cells will prevail in a given spatial domain if one watches the cell population dynamics for a sufficiently long interval because of its lower degree of contact inhibition (or smaller exclusion diameters).

The intrinsic randomness of biological phenomena justifies the use of a stochastic approach for their investigation. At intracellular level randomness is caused by chemical reactants being present in low copy numbers and their heterogeneous distribution inside the cell [4]. For example, random fluctuations have been widely observed in gene expression of either prokaryotic and eukaryotic cells by means of fluorescence techniques [5–20]. Despite randomness inevitability different gene regulatory strategies may give raise to different noise behaviors. For example, a self-repressing gene will present a higher precision on controlling the amounts of its products [1, 21–24]. Alternatively, external regulation has been noticed as a gene regulatory strategy resulting in higher noise [5, 25, 26]. Those results suggest self-repression as the unique mechanism controlling gene expression when high precision is necessary, as it is the case during development. However, recent results show that external regulation may be sufficient to generate the required spatial precision for gene products stripes formation along the anterior-posterior axis of a *D. melanogaster* embryo [2, 27, 28].

Indeed, developmental processes require a high precision on the control of the amounts of specific gene products such that they are present at proper positions and times during the life of an organism. That fact may induce a perception that noise is always detrimental to the cell. Such a premise is not always true. For instance, individual cells increase their survival chances under stress conditions by means of noise in gene expression with the consequential generation of phenotype diversification [29–32]. Normally behaving tissues are characterized by well organized cellular structures along space and time. That achieved by means of homeostatic mechanisms controlling cell densities in tissues. However, molecular fluctuations may affect cell genetics, induce uncontrolled expression of cell proliferation related genes, and induce the appearance of carcinoma *in situ*. The latter generates spatially less organized cell structures in tissues, breaks homeostatic behavior, and provides conditions for invasive cell phenotype to appear. That is a manifestation of stochasticity being a beneficial trait for cancer cells (at individual level) even though at an organismic level the noise has lethal effects after a while.

Therefore, an important challenge for cancer biologists is to determine mechanisms underpinning the progression of a carcinoma *in situ*, and how those cells become prevalent within a region for a sufficiently long interval such that invasive phenotype start appearing. One important mechanism necessary for the prevalence of the tumor cells is loss of contact inhibition [33, 34]. Contact inhibition of proliferation in culture experiments is associated with the ability of cells to maintain their density in a given tissue at optimal values [35, 36]. That causes cancer cells to keep proliferating in culture experiments even when confluence is reached [33]. In contrast, it has been shown that hypersensitivity to contact inhibition in fibroblasts of naked mole-rats is a mechanism that stops proliferation at lower cell densities in culture experiments. That is caused by the interplay between p16 and p27 cyclin-dependent kinase inhibitors which stops proliferation at lower densities than when p16 gene is not expressing such that cell densities reach that level observed in mouse [37]. Those experimental results suggest the necessity of finding a quantitative description of the intensity of contact inhibition in normal or cancer cells to enhance our capability of predicting or describing carcinoma *in situ* growth.

The next sections are devoted to give an overview of the three applications discussed above and to present some research perspectives. We start discussing the stochastic model for regulation of gene expression. We discuss the chemical kinetics that enable the self-repressing gene to be expressed at lower noise regimes. Furthermore, we present our results in the context of development of *D. melanogaster* embryos which indicates the possibility of using of this model to approach complex organisms. Then we discuss our cell level approach to quantify the degree of contact inhibition between two cells as an exclusion diameter. Lower degrees of contact inhibition are quantified by smaller exclusion diameters and that is applied on the description of a co-culture experiment with interacting melanoma and keratinocytes. We present some possibilities for future investigations at last section.

## II. RANDOM FLUCTUATIONS IN GENE EXPRESSION

Randomness in gene expression has been measured in terms of the Fano factor, denoted by 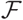, which is defined as the ratio of the variance to the average. We denote the number of gene products by *n* (the number of proteins or mRNA’s) and the Fano factor is written as:

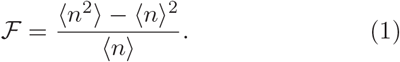

The Fano factor provides a measure to compare a probability distribution with the Poissonian distribution. The Poissonian distribution has 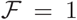 while 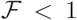 characterizes a sub-Poissonian distribution (a distribution that is thinner than the Poissonian). The a super-Poissonian distribution has 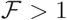 and is more spreaded than the Poissonian. Determining the probability distribution governing a the gene products number is important because it provides some hints on the regulatory strategy of the gene. For example, a constitutive gene will have *n* being a random variable governed by a Poisson distribution, a sub-Poisson distribution governs the self-repressing gene [1], while super-Poissonian distributions might indicate a positive feedback (governed by a bimodal distribution) or bursty expression (governed by gamma or negative binomial distribution).

The above analysis is completed in the scope of regulation of gene expression modeled by means of a binary promoter that is ON or OFF. When the promoter is ON there is synthesis of gene products at rate *k* while no synthesis happens in the OFF state. The gene products decay at a rate *ρ*. The rate of promoter switching from OFF to ON state is denoted by *f* while the opposite transition happens at rate *h*_1_ (for the self-repressing gene) or rate *h*_2_ (for the externally regulating gene). Fig. 1 presents a cartoon of our simplified model for regulation of gene expression.

**FIG. 1:**
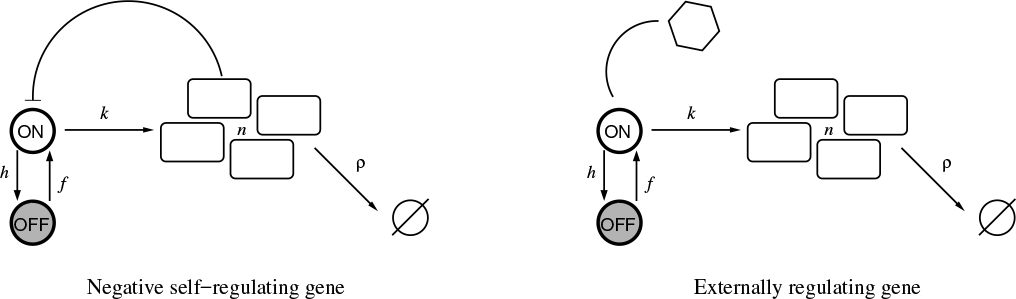
The left cartoon is a representation of a self-repressing gene while the right cartoon represents an externally regulated gene.

The scheme presented in Fig. (1) correspond to the set of effective chemical reactions presented below. The left hand side equations correspond to the self-repressing gene while externally regulated gene effective reactions are presented on the right. For the SRG we denote a protein by 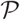. The regulatory region of the gene is denoted by 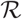 and the gene state is determined by the binding of 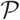 to the regulatory region. The regulatory protein of the externally regulated gene is denoted by 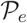. The symbols for reaction rates appear on top of arrows indicating the reactants and products of the effective reactions.

Self-repressing gene

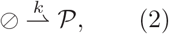

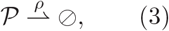

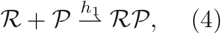

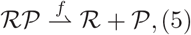

Externally regulating gene

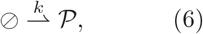

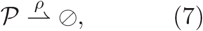

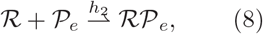

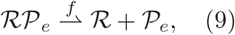

The Eqs. (2) and (6) are indicating protein synthesis while protein degradation is indicated by Eqs. (3) and (7). The gene switching from ON to OFF state is indicated by Eqs. (4) and (8) while the opposite transition is presented at Eqs. (5) and (9). The system of effective reactions presented here is very simplified in comparison with the complexity of gene regulation and gene expression in mammals. However, such a simplification is highly necessary for establishing a quantitative description based on exactly solvable models.

A stochastic model for the binary gene with the probabilities of finding the gene at the ON (or OFF) state when *n* gene products are found inside the cell being denoted by *α_n_* (or *β_n_*) have been presented elsewhere [38, 40]. Hence, the state of the system is determined by two random variables (*m, n*), with *m* =(ON,OFF) and *n* being a non-negative integer. Those probabilities can be computed for a specific stochastic process determining their evolution which can be done by means of continuous time Markov processes also known as master equations. The master equations are characterized by a combination of the individual transformations that change the state of the system. The left-hand side of a master equation has the rate of change of the probability for the system being in a given state while the right-hand side has the processes that are causing the probabilities to change. At the left-hand side of the master equation, a positive contribution means that the transformation brings the system to current state while transformations taking the system from the current state towards a different one give negative contributions.

The master equations governing the dynamics of the probabilities (*α_n_*,*β_n_*) are written below. We provide an interpretation for the first term on the right-hand side and the remaining can be interpreted in the same framework. The term proportional to *k* has a positive component *α*_*n*−1_ and *α_n_* as a negative component. The former means that if the state of the system is (ON,*n* − 1) and there is a synthesis of a gene product the system reaches state (ON,*n*) while the second means that a synthesis takes the system from current state (ON,*n*) towards state (ON,*n* + 1). The master equation is written as

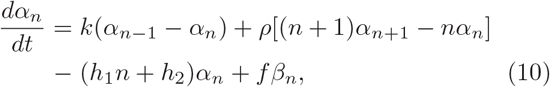

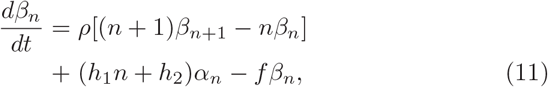

where the self-repressing gene is modeled considering *h*_1_ ≠ 0 and *h*_2_ = 0, such that the switching rate from ON to OFF state depends on *n*. Note that in the self-repressing gene the ON to OFF switching rate has linear dependence on n. The contrary condition, *h*_1_ = 0 and *h*_2_ ≠ 0, results in a model for the externally regulated gene. The solution to the Eqs. 10 and 11 have been obtained exactly for the self-repressing gene [38, 39] and the externally regulated gene [40, 41]. For the two rates *h*_1_ and *h*_2_ being non-null one has a model for a gene that is both self-repressing and externally regulated. That will not be approached here because it is a subject for additional research.

A striking feature of biological organisms is their capability of regulation that ensures that a given gene will be expressed in proper quantities with spatial and temporal precision. Hence, though variation in gene amounts is observed these fluctuations are within specific ranges in normally behaving biological systems. An important question is to find regulatory strategies underpinning such a precision such that a classification of regulatory strategies and their biological function would emerge. For example, there are experiments demonstrating that self-repressing genes are responsible for reducing random fluctuations in gene expression [21, 22, 24]. Indeed, it has been shown that self-repression induces noise reduction such that one may obtain sub-Fano probability distributions [23]. However, the mechanisms enabling such a reduction in noise were not clear. That has appeared later by writing the Fano factor as

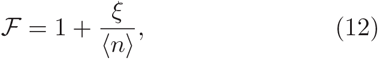

where ξ is the covariance between the variables (*m, n*) with the values of *m* (ON,OFF) being represented by corresponding synthesis rates (*k/ρ*, 0) such that ξ can be computed. The self-repressing gene has the possibility of ξ < 0 and sub-Fano probability distributions will occur [1]. Those regimes will happen when the prevailing process of the system is the gene switching between the ON and OFF states while few synthesis or degradation of gene products happen during a given time interval.

The left graph of Fig. 2 shows the Fano factor for a self-repressing gene in the sub-Fano regime. Note the existence of the possibility of finding arbitrarily low values for 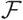 when 〈*n*〉 = 1. That corresponds to a kinetics with the regulatory protein having a high affinity the regulatory region controling the expression of the gene. At that regime, once a regulatory protein is released from the DNA and the gene becomes ON, an available protein rapidly binds to the DNA and the gene switches back to the OFF state.

The cartoon that we are considering for gene regulation may seem a strong simplification of the the whole picture in metazoans. However we may use that approach for description of eukaryotes under specific assumptions. For example, during its early developmental stages *D. melanogaster* embryos are characterized by a syncytium such that the cells only have their nuclei. That enables us to apply the gene transcription model for an externally regulating gene and use it as a first step to understand how spatial noise appears because of noise on amounts of gene products.

**FIG. 2:**
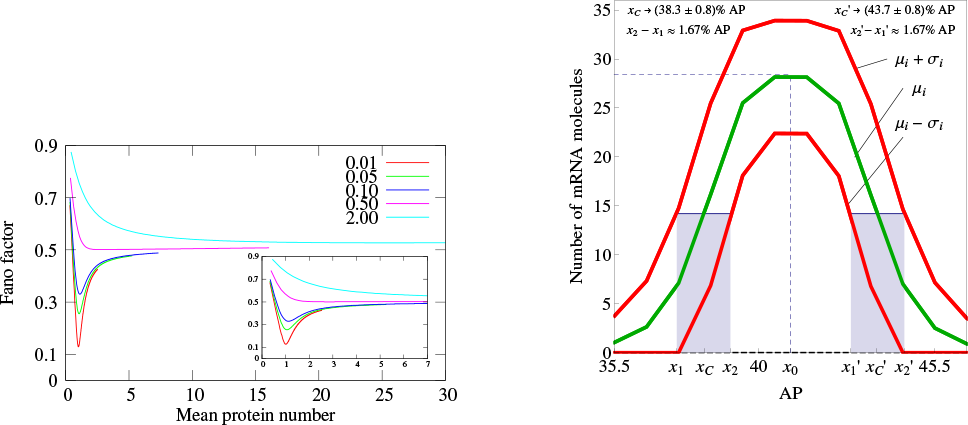
Left graph. Fano factor versus average protein number for the self-repressing gene. The value of *a* is fixed as 500. The values of *b^s^* are indicated within the graph while we have varied the value of *z*_0_.*Right graph*. Spatial profile of mRNA’s average amounts along AP axis of a *D. melanogaster* embryo. We also include the fluctuation on the position of the borders of the peak of expression accordingly with the standard deviation of *n* at each nucleus along AP axis. The position of the borders are computed at the point where 〈*n*〉 is half of its maximal value at position 41.5 % of the embryo length.

Indeed, we carried out that approach to model the *even-skipped (eve)* gene which is important for the formation of spatial protein concentration patterns that in the adult organism will determine specific segments with different functions [2]. *eve* mRNA’s spatial pattern is characterized by a Gaussian profile at the onset of gastrulation (see right graph of Fig. 2). To apply our model we assumed a one-dimensional lattice where each node has a single copy of *eve*. The lattice represents the AP axis of the embryo and theoretical values for 〈*n*〉 at each node of the lattice were compared with observed values for the fluorescence of mRNA of *eve* as obtained experimentally [42]. At this stage the challenge was to propose a method for converting the detected intensity of immunofluorescence into mRNA numbers. Then we compared the two spatial patterns at the onset of gastrulation (theoretical and experimental) and obtained a good agreement. The second stage was to compute the values of 〈*n*〉 ± *σ* along the whole lattice, where *σ* indicates the standard deviation on *n*. Then we compared the position of the borders of the domains and their fluctuations with experimental data [27]. That showed that the theoretical fluctuations were of the same order as of the experimental ones. That is an unexpected result since it indicates that the spatial precision required during mRNA numbers pattern formation in developing embryos can be achieved with a gene transcription regulatory strategy that is not the most precise on controlling transcripts numbers.

## III. CELL LEVEL MODELS

There are strong experimental evidence that loss of contact inhibition is a key element underlying the capability of tumors to grow and prevail within a given tissue (forming the *carcinoma in situ*) [33–36] or that hypersensivity to contact inhibition prevents such a prevalence [37]. Those roles indicate the necessity of understanding quantitatively (and geometrically) how contact inhibition affects the dynamics of cell proliferation and the spatial patterns that they form in a given tissue. Such an approach is going to be useful for the understanding of early cancer development and future design of techniques for early cancer diagnosis and treatment. To approach that problem we have recently proposed a co-culture experiment combining keratinocytes (HaCaT or normal) and melanoma (SK-MEL-147 or cancer) cells [3]. We considered an initial configuration of 10:1 (keratinocytes:melanoma cells) and evaluated the cell density daily until confluence was reached. The initial configuration was composed by well-mixed populations of keratinocytes and melanoma cells. At confluence we observed spatial patterns with spreaded normal cells surrounding melanoma cells clusters (see Fig. 3.G). In that experiment the growth of the two sub-populations of cells was tracked and their individual curves are easily adjusted by Gompertz logistic-like curves [43] (see fig. 3.B). Gompertz curves are characterized by a sigmoidal shape such that the population density has a slow growth rate at early phase. That growth rate increases until it reaches a maximal value and starts diminishing while cell population density reaches its maximal value asymptotically. The density growth rate is proportional to a constant which is interpreted as the inverse of the cell division time during the early phase of culture experiments when the growth is still approximately exponential. The fitting has shown that the growth rate of both cell types had same value while the final proportions of the two populations has diminished from 10:1 to ≈ 4: 1 (Fig. 3.C shows the temporal evolution of this ratio). That result shows a limitation for using Gompertz-like curves to describe experimental results when distinct cell types interact in the same environment. Particularly, those models have been developed in the context of predator-prey interacting in an environment with finite resources availability [44]. We did not found in literature any instance of co-culture of melanoma and normal cells interacting similarly to a predator-prey system. Hence, a different approach to describe our system quantitatively was required.

To obtain a quantitative description of our model we consider the a variation of the Widom-Rowlinson model [45–47]. A cartoon of our model is presented in Fig. 4.A where the tissue is represented as a two-dimensional grid of size *L*×*L*. Melanoma (or normal) cells, indicated in red (or blue), can occupy the grid’s vertices with the distance between two cells being the smallest amount of edges connecting their vertices. The latter enable us to define contact inhibition by means of an exclusion diameter around the cell as it is represented by the shadowed areas around the red circles of Fig. 4.A. The vertices within the purple regions cannot be occupied by the melanomas while the normal cells cannot occupy the vertices within the red shadowed areas. The exclusion diameters are reciprocal and notice that the melanoma cells occupy their immediate neighbors. In our model the cell type *i* (*i* = 1 or *i* = 2, respectively, for melanoma or normal) undergo division (with rate *α_i_*), quiescence (with rate *σ_i_*), death (with rate *ρ_i_*) and migration (with rate *δ_i_*) with specific rates.

**FIG. 3:**
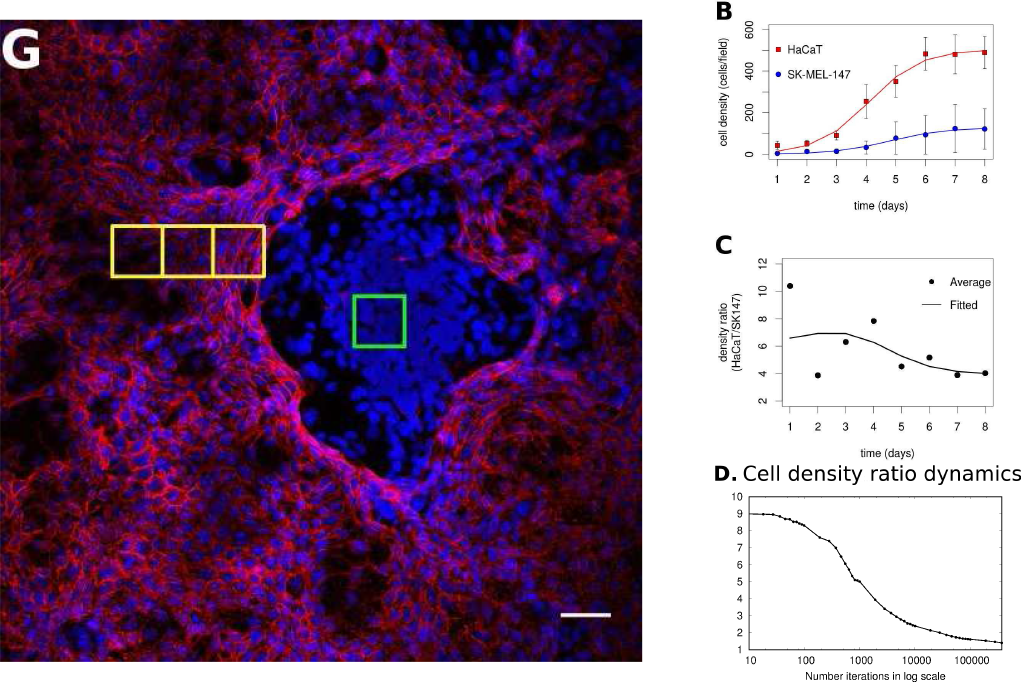
Graph G. shows a representative configuration of the co-culture experiment at the confluence regime. Graph B. gives the evolution of the individual populations along until confluence is reached and their fitting by a sigmoidal curve. Graph C. the experimental ratio of melanoma to normal cells along time is shown. Graph D. the simulation of the ratio of melanoma to normal cells along time is shown.

In Ref. [3] the dynamics of the model is established by means of a Monte Carlo Markov Chain which is briefly described below. A vertex *x* of the grid is selected with probability *L*^−2^ and its state is verified. (*1.*) For the vertex being **occupied** by the *i*-th cell type: the cell is quiescent, *i.e. x* remains occupied, with probability *α_i_*/*Q*; there is a probability 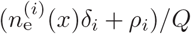 for vertex *x* turn empty, where 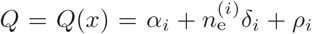. The vertex *x* turns empty because of cell *death* - with probability *ρ_i_*/*Q* - or *migration* - with probability 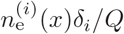. In case of *migration* the cell arrive at any vertex at distance *D*(*i,i*) that satisfies the admissibility rule. The number of vertices around *x* that can receive the cell is indicated by 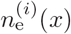 and 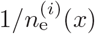 is the probability for one of these vertices receive the migrating cell. (*2.*) For the vertex *x* being **empty**: it remains empty with probability (*ρ_1_* + *ρ_2_*)/*R* [^1^ Although that probability might be defined by means of a different rate our choice avoided introducing more parameters to our simulations.]; the vertex *x* may become occupied by the *i*-th cell type with probability *n_i_* (*x*)(*α_i_* + *δ_i_*)𝕀(*i*)/*R*. The occupation of *x* may happen because of a cell *division* - having probability *n_i_*(*x*)*α_i_*𝕀(*i*)/*R* - or *migration* - having probability *n_i_*(*x*)*δ_i_*𝕀(*i*)/*R*, where 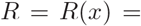 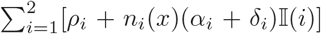. The amount of allowed nearest vertices occupied by the *i*-th cell type around *x* is denoted by *n_i_* (*x*). A migrating cell has a probability 1/*n_i_* (*x*) to be chosen and, after migration, the vertex from which the cell comes from is turned empty. The symbol 𝕀(*i*) is equal to 1 or 0 when the vertex *x*, respectively, can or cannot receive the *i*-th cell type accordingly with admissibility rule. Note that *x* remains empty with probability one for *R* = *ρ*_1_ + *ρ*_2_.

**FIG. 4:**
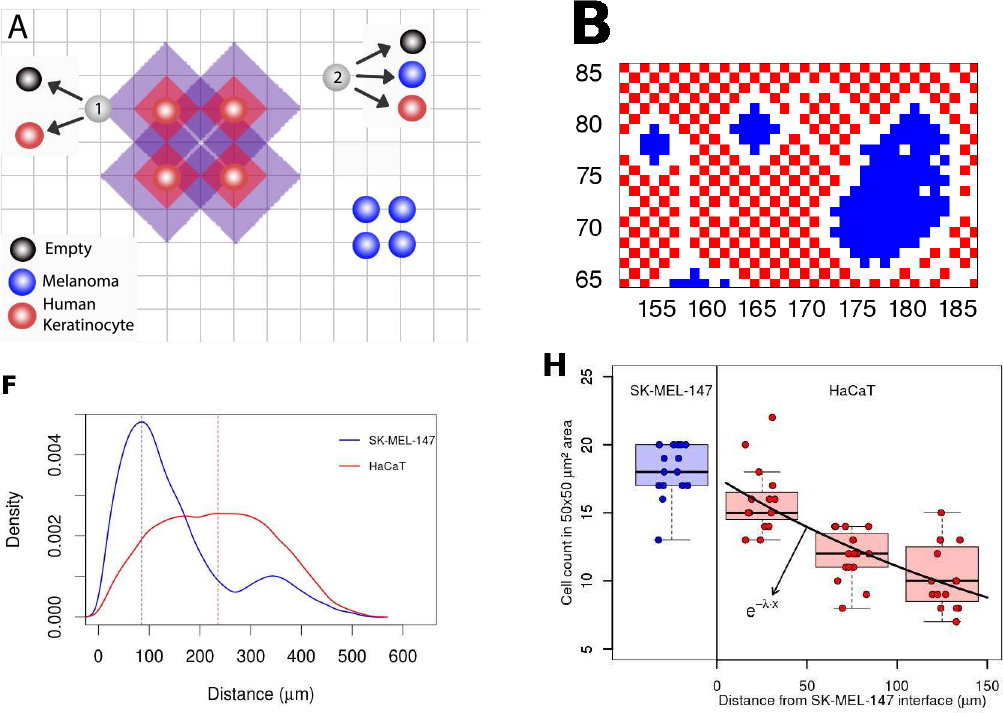
Graph A. gives a cartoon of our model for proliferation under different allelophylic degrees. Graph B. is a spatial configuration achieved in our simulations at the co-culture confluence regime. Graph F. gives the experimental cell-to-cell distance distribution (the blue (red) curve indicates cancer (normal) cells histogram). Graph H. shows how normal-to-normal cell distance as they are further away from the interface with the melanoma clusters.

Fig. 3.D shows the results of our simulations for the ratio between the densities of normal to tumor cells. We show that that ratio follows the same pattern as observed in our experiments. Furthermore, the densities of the two sub-populations also follow the same pattern (as it may be verified in Fig. [3]) such that the kinematics obtained with our model adjusts to that observed experimentally. Hence, it is fair to conclude that our model is a strong candidate to quantify how different degrees of contact inhibittion affects the dynamics of cell proliferation in co-culture experiments, and, maybe, in tissues.

Indeed, fig. 4.B shows a spatial configuration that we obtain after simulating our model. The blue circles form clusters that are surrounded by the red ones. This pattern is similar to the ones we have observed in our coculture experiments, fig. 3.G. Furthermore, we also evaluated the typical cell to cell distances observed in our experimental images. Fig. 4.F shows that at confluence regime normal to normal cells typical distance is twice that between cancer cells. That reinforces of our geometrical interpretation of contact inhibition by means of an exclusion diameter and shows the cancer cells as *allelophylic* (*allelo*, the other; *phylia* affinity). In our study we also found the correspondence between the spatial scales of the simulation and the experimental ones. That has enabled us to compare the melanoma clusters characteristics observed experimentally with those of the simulations [3].

## IV. PERSPECTIVES

Our results raises perspectives for further research, and a non-exhaustive set of possibilities is discussed below. The use of a stochastic binary model for gene expression on the description of the formation of stripe two of *eve* along the AP axis of the *fruitfly* embryo indicates the necessity of understanding the effects of fluctuations on molecular numbers on the spatial organization of cells. On the other hand, the use of model for a self-repressing gene can be used in cancer context to investigate the behavior of BACH1 production under influence of a biometallic compound such as heme [48]. BACH1 is a transcription factor with negative self-regulation [48] found overexpressed in triple-negative breast cancer cells. Since it has been demonstrated its role as a metastasis promoter, to model how metastic breast cencer cells keep it at higher levels is relevant to the development of new therapuetic approaches to this condition. Heme accelerates BACH1 decay and the model may help on the design of treatement strategies that also reduce both the expected amounts of BACH1 within the cell and their fluctuations such that phonotype heterogeneity of tumor cells is reduced. That reduction has the potential of increasing the cancer treatments effectiveness and may establish guidelines to reduce the invasive and metastatic capabilities in cancer cells. Under a more theoretical perspective, we may also consider the investigation of the meaning of the symmetries of the stochastic binary models [23, 26] aiming to model two interacting genes.

Another possibility of cell levels models is to propose Markov chains to approach tumor heterogeneity at the phenotype level. The tumor progression can yield changes in its architecture that lead tumor cells to die or to develop invasive phenotypes because of scarcity of space and resources [49–52]. Additionally, environmental cues may regulate expression of transcription factors that regulate the internal cell dynamics [53–56] from proliferative towards invasive. Those facts suggest the proposition of a cell level model for tumor progression having two cell phenotypes, with the invasive being originatedfrom the proliferative, and population dependent transition rates as an effective homeostatic mechanism. On more specific possibilities, the use of the stochastic model for contact inhibition may also be extended to the condition when there are three or more interacting cell types. In that case, we may start with a three cell states, accounting for keratinocytes, melanocytes, and melanomas. Here, the melanomas would result from a modification of melanocytes and we may use those results to investigate the conditions for the progression of a melanoma *in situ* from a normal configuration. The cell level approaches have investigated the two dimensional grids for a description of culture experiments. Our next step is to construct three dimensional grids to enable us to describe *in vivo* experiments and, hence, obtain a richer picture about carcinogenesis. One natural challenge of such an approach is to establish the grid’s topology such that different first neighbors rules would come up. Those new simulations would enable us to develop new imaging analysis tools that will be useful for a quantitative characterization of the spatial patterns that are generated at different stages of carcinogenesis. Those characterization may motivate the development of automated tools for tissue characterization by patologists.

Fig. 3.G shows four yellow squares, one within a melanoma cluster and the remaining three sequentially located from the interface of normal and melanoma cells domains. Inspection is sufficient to verify that normal cells nearby melanomas are closer to each other than those at a longer distance from the cluster. That pattern repeats in our experiments and a graph of the densities of keratinocytes at different distances is shown in Fig. 4.H. Box-plots for melanoma-to-melanoma (or normal-to-normal) cell densities at different distances are presented. We show that the cell-to-cell separation falls exponentially which suggests the existence of a molecular mechanisms determining that separation which concentration is dependent of the presence of the melanoma cells. That points out towards the necessity of combining our two approaches for cancer biology at both molecular and cellular levels. That would be useful to determine how molecular level fluctuations give raise to cancer heterogeneity.

Our investigations on molecular mechanisms of carcinogenesis may also have implications for the analysis of low dose and low dose rates rates random effects [57, 58]. It is estimated that radiation therapy accounts for 50% of cancer treatment cases and that application may be playing a role on late appearence of tumors. Hence, understanding how low dose and low dose rates of ionizing radiation are affecting cells may be an important scientific problem with clinic implications. At those regimes one expects the effects of ionizing radiation to be stochastic such that the it is natural to employ an approach based on probabilistic theory. Initial attempts for stochastic modeling of biological effects are based on target theory [59] while more detailed deterministic models have been proposed recently to account for DNA repair mechanisms of mammal cells [60]. For the latter, we will employ the Langevin technique to evaluate randomness in deterministic models. On a different research direction, one may notice that 90% of radiation treatments use radiofrequency-driven linear accelerators of electrons (RF-Linac). These RF sources are not very precise and may lead radiation to hit both the tumor and neighboring cells. Hence, there are continuous efforts for finding new radiation sources including laser accelerated electrons for the generation of tunable and quasi-energetic x-ray sources [61, 62]. The development of that technology relies hardly on computational simulations of the interaction of laser plasma interactions devoted to the optimization of the generation of x-rays through electron acceleration by lasers. Those simulations may guide experimental designes which would result on the proper generation of x-rays for clinical purposes.

## ACKNOWLEDGMENTS

This work has been supported by CAPES Process n. 88881.062174/2014-01 and have used computational resources of Department of Information Technology of University of São Paulo. AUS and MFSV have been supported by PET program of Ministry of Education of Brazil. MYS, ASQ, MCCM have been supported by CAPES Programa de Pós-graduação em Oncologia of Faculdade de Medicina, Universidade de São Paulo.

## AUTHOR CONTRIBUTIONS

AUS, MFV, MYS, WWL, ASQ, MCCM, collected data, wrote the manuscript and approved the final version of the manuscript to be published. AFR designed the study, collected data, interpreted the results, wrote the manuscript and approved the final version of the manuscript to be published.

